# Development of the overlapping network modules in the human brain

**DOI:** 10.1101/2024.05.03.592316

**Authors:** Tianyuan Lei, Xuhong Liao, Xinyuan Liang, Lianglong Sun, Mingrui Xia, Yunman Xia, Tengda Zhao, Xiaodan Chen, Weiwei Men, Yanpei Wang, Leilei Ma, Ningyu Liu, Jing Lu, Gai Zhao, Yuyin Ding, Yao Deng, Jiali Wang, Rui Chen, Haibo Zhang, Shuping Tan, Jia-Hong Gao, Shaozheng Qin, Sha Tao, Qi Dong, Yong He

**Author notes:** **Corresponding authors:** (XL), (YH).

## Abstract

Developmental connectomic studies have shown that the modular organization of functional networks in the human brain undergoes substantial reorganization with age to support cognitive growth. However, these studies implicitly assume that each brain region belongs to one and only one specific network module, ignoring the potential spatial overlap between functional modules. How the overlapping functional modular architecture develops and whether this development is related to structural signatures remain unknown. Using longitudinal multimodal structural, functional, and diffusion MRI data from 305 children (aged 6–14 years), we investigated the development of the overlapping modular architecture of functional networks, and further explored their structural associations. Specifically, an edge-centric network model was used to identify the overlapping functional modules, and the nodal overlap in module affiliations was quantified using the entropy measure. We showed a remarkable regional inhomogeneity in module overlap in children, with higher entropy in the ventral attention, somatomotor, and subcortical networks and lower entropy in the visual and default-mode networks. Furthermore, the overlapping modules developed in a linear, spatially dissociable manner from childhood to adolescence, with significantly reduced entropy in the prefrontal cortex and putamen and increased entropy in the parietal lobules. Personalized overlapping modular patterns capture individual brain maturity as characterized by brain age. Finally, the overlapping functional modules can be significantly predicted by integrating gray matter morphology and white matter network properties. Our findings highlight the maturation of overlapping network modules and their structural substrates, thereby advancing our understanding of the principles of connectome development.

## Introduction

The period of childhood and adolescence is a stage of transition from infancy to adulthood, which is critical for the maturity and improvement of motor, cognitive, emotional, and social functions [1, 2]. During this period, the brain undergoes progressive and regressive maturation in its microscopic and macroscopic anatomy, such as increased myelination [3, 4], synaptic pruning [4–6], and cortical thinning [4, 7–9]. From the perspective of function, remarkable reconfigurations have also been observed for task-evoked regional activity [10, 11] and task-free spontaneous activity [12]. These structural and functional changes are deemed to lay the foundation for improvements in children’s cognitive and behavioral performance [10, 11, 13].

Notably, the periods of childhood and adolescence are also critical windows for the onset of many psychiatric disorders [14]. Exploring brain developmental principles in children and adolescents offers novel insights into the neural mechanism underlying not only cognitive and behavioral growth but also neurodevelopmental disorders.

Over the past two decades, neuroimaging-based connectomics has provided a valuable framework for investigating the developmental principles of brain function [15–17]. The functional modular structure, which is characterized by dense within-module connections and sparse between-module connections, has attracted great attention [18–21]. The modular structure of the brain is particularly important for global network communication since it can facilitate efficient information segregation and integration with low wiring costs [20, 22]. Several studies have reported age-related changes in modular organization with development [15, 16, 23–25].

Specifically, functional modular organization is already present in fetuses [26] and neonates [27, 28]; in this configuration, the modules in the primary cortex show an adult-like topography, and the modules in the association cortex are far from mature. The modular architecture further undergoes an elaborate reconfiguration from childhood to adulthood [15, 16, 23, 24]. Its spatial layout shifts from an anatomical proximity to a spatially distributed and functionally related configuration [25]. These developmental changes have been found to be related to the development of individual cognition and behavior, such as cognitive control [24] and general cognitive function [23].

Despite this substantial progress, previous connectome development studies have focused primarily on modular structure without spatial overlap (i.e., hard assignment), implicitly assuming that each brain node belongs to one and only one functional module. This hypothesis might be problematic because the modular structures of real-world networks, such as cooperation networks in social systems and protein networks in nature, generally show overlapping properties [29, 30]. This modular overlapping framework in complex networks provides important insight into the potential diverse functional roles of nodes in the network. Several recent studies have also reported overlapping modular organization in brain functional networks in adults [31–35], indicating that brain regions are not limited to a specific module. Specifically, the overlap between functional modules is spatially heterogeneous [32–35], with higher overlapping regions being crucial for intermodule communication [36] and global network efficiency [33]. The presence of overlapping modules is also related to spatial heterogeneity in functional diversity and neurocognitive flexibility [32, 33, 37]. However, the network growth principle of the overlapping modular organization in the functional connectome and its association with structural brain signatures remain unknown.

To fill these knowledge gaps, we investigated the development of the overlapping modular architecture of functional connectomes during childhood and adolescence and examined their associations with structural brain signatures using a large longitudinal multimodal neuroimaging dataset from 305 typically developing children (aged 6–14 years, 491 scans in total) [38, 39]. Specifically, we used resting-state fMRI (rsfMRI) data to investigate the development of the overlapping modules and structural and diffusion MRI data to reveal the underlying structural substrates of the overlapping modules. In the present study, we used an edge-centric module detection approach [40] to identify the overlapping functional modular architecture for each rsfMRI scan of each participant and further quantified the degree of overlap of each brain node in the module affiliations. We then examined development in the overlapping level of child brain networks at the global, system and nodal levels, and further assessed whether the spatial pattern of the modular overlap can predict brain maturity. Finally, we investigated the potential structural substrates involved in the development of overlapping functional modules.

## Results

We leveraged longitudinal rsfMRI data from 305 children (aged 6-14 y, 491 scans), including three repeated scans from 47 children, two repeated scans from 92 children, and one scan from 166 children (Figure 1A). For comparison purposes, we also included cross-sectional rsfMRI data from a group of healthy adults (n = 61, aged 18-29 y). Both children and adults were scanned using the same scanner with identical protocols. All MR images used here underwent strict quality control (for details, see Materials and Methods). We identified the overlapping modular architecture in individual- and group-level functional networks using an edge-centric modular detection algorithm [40] (Figure 1B). Briefly, we first constructed a traditional functional network comprising nodal regions and interregional connections (i.e., edges). Then, we constructed a corresponding weighted edge-based brain graph that represented the similarity of connectivity profiles between edges (see Materials and Methods). Finally, we identified module affiliations of each nodal region according to the module assignments of its edges in the corresponding edge graph. A measure of entropy was used to measure the extent of modular overlap for each node by quantifying the distribution of module affiliations of the edges attached to this node (Figure 1B).

**Figure 1.**
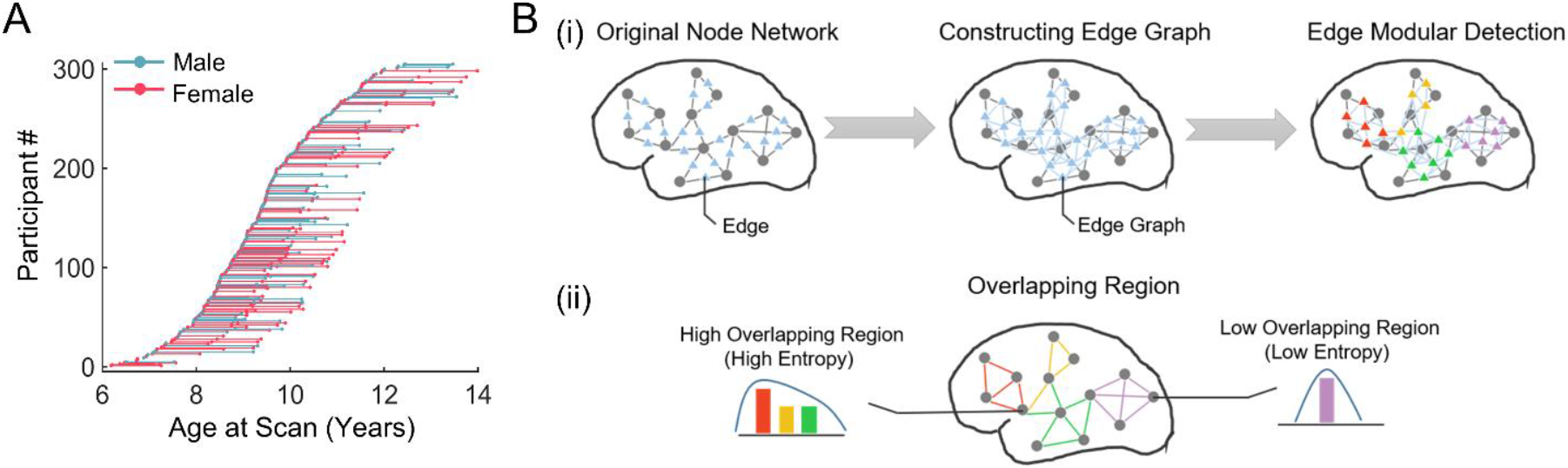
Data information and schematic diagram of the overlapping modular architecture based on edge-centric module detection. **(A)** Age distribution of longitudinal rsfMRI scans of children. **(B)** (i) Left: traditional brain functional connectivity network. In this network, each node denotes a brain region of interest, and each link denotes the interregional functional connectivity. Middle: edge graph corresponding to a given functional network. In this graph, each node denotes an edge in the functional network, and each link is defined as the similarity between edges in the connectivity profiles. Right: edge-centric module detection. Each edge is assigned to a specific module based on the Louvain algorithm [41]. (ii) Definition of regional module overlap. Each nodal region was assigned to one or more modules due to the diverse module affiliations of its edges. We employed a measure of entropy to quantify the extent of module overlap of each brain region by measuring the distribution of the module affiliations of its edges.

### Spatial topography of the overlapping functional modules in children and adults

We first identified the overlapping functional modules in healthy young adults, which serves as a reference for exploring the development of the overlapping modules in children. We found seven modules in the weighted edge-based brain graph of the adult group (Figure 2A) and further showed the corresponding topographic distribution of each module (Figures 2B and 2C). These functional modules showed substantial spatial overlap, as characterized by 73% of the nodal regions belonging to two or more modules (Figure 2B). Module I was mainly located in the medial and lateral prefrontal and parietal cortex, and lateral temporal cortex; module II was mainly located in the primary motor and somatosensory cortices; module III was mainly located in the insula, supramarginal gyrus, superior temporal gyrus, and paracentral lobule; module IV was mainly located in the cingulate gyrus and the subcortical area; module V was located in the visual cortex and the superior parietal lobule; module VI was located in the middle frontal gyrus and superior parietal lobule; and module VII was primarily located in the temporal pole, hippocampus, and amygdala (Figure 2C).

**Figure 2.**
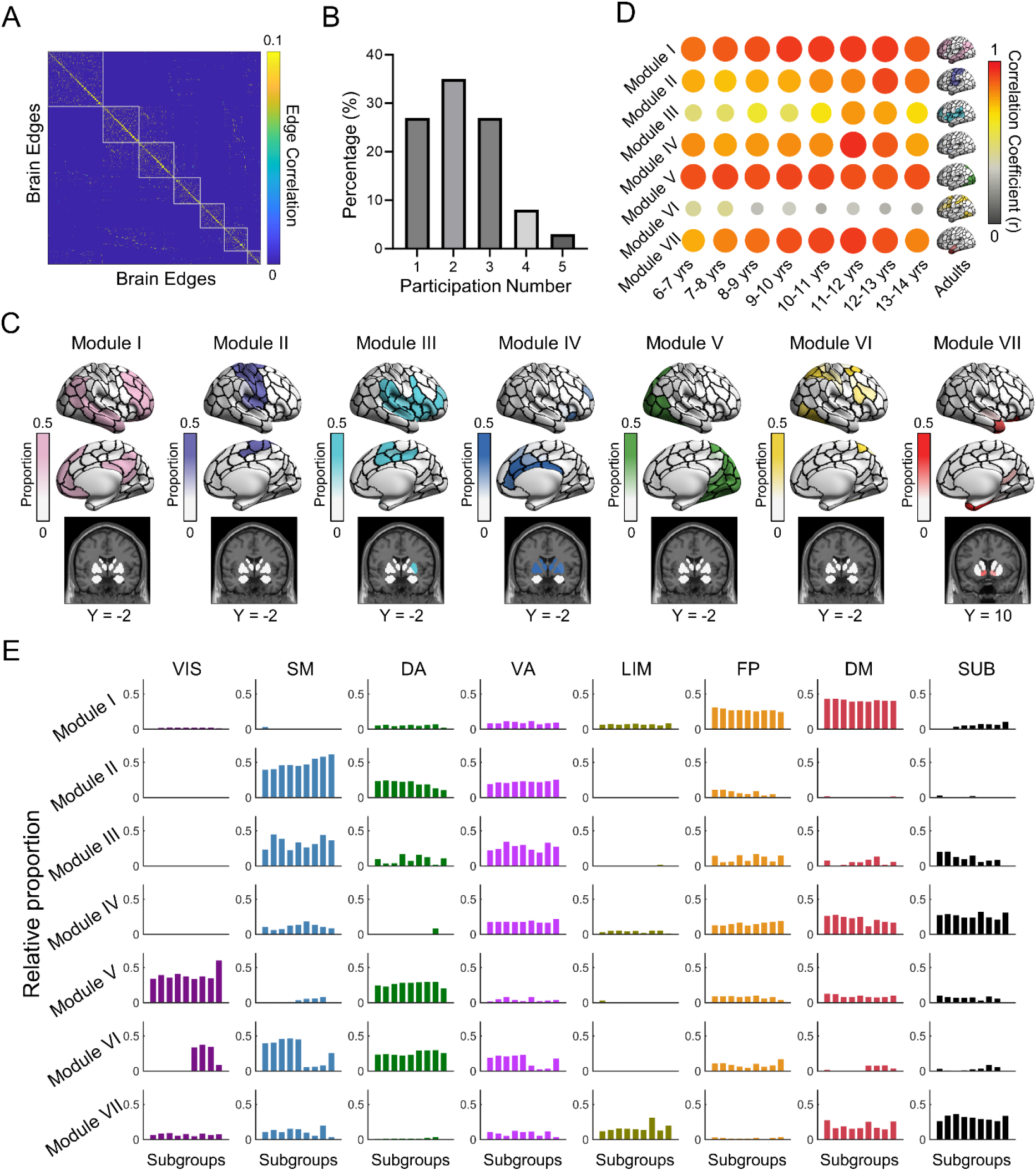
Overlapping modular architecture of the brain functional network in the adult cohort and the child subgroups. **(A)** Modular organization in a weighted edge graph. The edges were sorted according to their module affiliations. Each element denotes the interedge similarity in their connectivity profiles. **(B)** Distribution of the number of nodal regions involving functional modules. Notably, 27% of the nodes belonged to one module, and 73% of the nodes belonged to two or more modules. **(C)** Topographic distribution of seven functional modules. For each module, nodal values represent the proportion of edges assigned to that module. **(D)** Spatial similarity of the functional module maps between the child subgroups and the adult group. The color and size of the dots represent the spatial similarity of functional modules between each child subgroup and the adult group. Larger dot sizes indicate higher spatial similarity. **(E)** System-dependent spatial distribution of functional modules. For each functional module of each subgroup, we calculated the percentage of nodes distributed in eight systems, including seven functional systems and the subcortical area. Given a prior system, the bar chart shows the percentage of nodes located in this system for eight child subgroups with a one-year interval and the adult cohort. In (C) and (D), cortical data were mapped on the brain surface using BrainNet Viewer software [46]. VIS, visual; SM, somatomotor; DA, dorsal attention; VA, ventral attention; LIM, limbic; FP, frontoparietal; DM, default-mode; SUB, subcortical; yrs, years.

We further divided all the children’s rsfMRI scans into eight subgroups with one-year intervals, and the adult group was set as the ninth subgroup for comparison. We identified the overlapping modular architecture for each subgroup of children based on the rsfMRI data. The modules in each child subgroup were matched with those in the adult subgroup. In general, most functional modules in the child subgroups showed high spatial similarity with those in the adult subgroup (Pearson’s correlation *r*s: mean ± SD = 0.75 ± 0.02, range: 0.25-0.95) (Figure 2D). Visual inspection suggested that the spatial similarity with the adult group increased gradually with age for all modules, except module VI. For each module, the spatial distribution of nodes among prior functional systems [42, 43] was largely consistent across all child subgroups and the adult cohort (Figures 2E). Module I mainly involved the default-mode and frontoparietal systems; module II mainly involved the somatomotor, ventral attention, and dorsal attention systems; module III mainly involved the somatomotor and ventral attention systems; module IV mainly involved the ventral attention, frontoparietal, and default-mode systems, and the subcortical area; module V mainly involved the visual and dorsal attention systems; module VI mainly involved the visual, somatomotor, dorsal attention, and ventral attention systems; and module VII mainly involved the limbic and default-mode systems and the subcortical area. Interestingly, we found that each module mainly comprised nodal regions that belong to the functional systems along the adjacent hierarchy [44, 45], regardless of the subgroup.

### Development of the overlapping functional modules during childhood and adolescence

We employed the mixed effect model [47, 48] to quantify the longitudinal changes in the overlapping modular structure during childhood and adolescence. At the global level, the number of modules in the edge-based brain graphs significantly decreased with age (linear model, *t* = - 2.09, *p* = 0.037), and the modularity tended to increase with age (linear model, *t* = 1.89, *p* = 0.059) (Figure 3A). At the regional level, the spatial topography of the nodal overlap (i.e., entropy) of every child subgroup was highly similar to that of the adult subgroup (Pearson’s correlation *r*s ranged from 0.74 to 0.91). Specifically, for each age subgroup, regions with higher levels of overlap were located mainly in the insula, supramarginal gyrus, inferior frontal gyrus, somatosensory cortex, anterior cingulate gyrus and subcortical area (e.g., putamen), and regions with lower levels of overlap were located mainly in the visual cortex, angular gyrus, and posterior cingulate gyrus (Figure 3B). Statistical analysis revealed that seven nodal regions showed significant linear changes with age (*pFDR corrected* < 0.0014, Figure 3C). These regions showed dissociable age-related changes, with significant increases mainly located in the superior and inferior parietal lobules and the lateral prefrontal cortex and significant decreases mainly located in the ventral and medial prefrontal cortex and the putamen.

**Figure 3.**
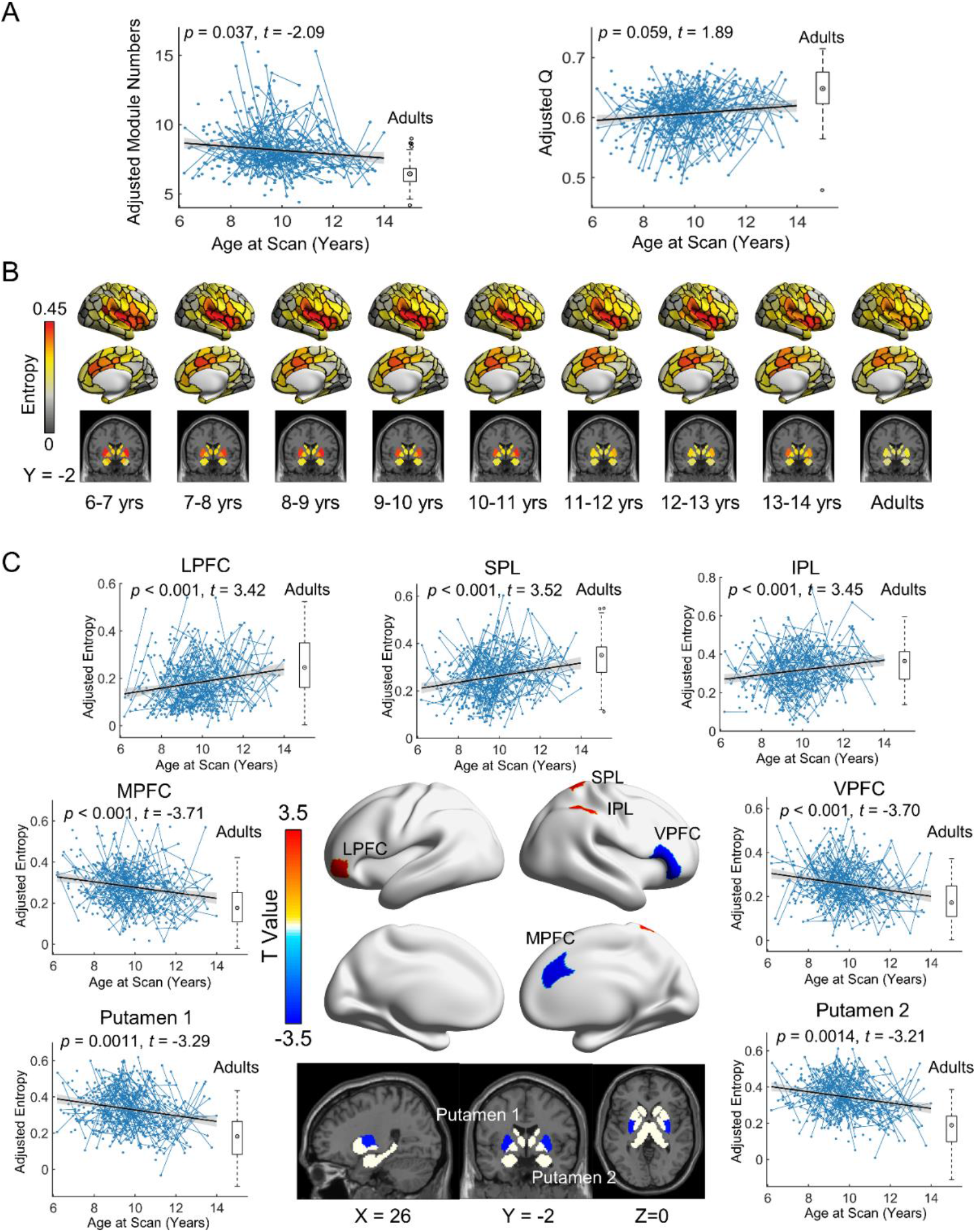
Longitudinal development of the overlapping functional modules at the global and nodal levels. **(A)** Left: Effect of age on the number of modules. Right: Age effect on modularity in edge-centric module detection. **(B)** Spatial patterns of functional module overlap across the brain for each child subgroup and for the adult group. **(C)** Spatial distribution of regions showing significant developmental changes in functional module overlap. Age effects are displayed in terms of *T* values (*pFDR corrected* < 0.0014). In (A) and (C), boxplots represent the distribution of the adult group for reference. The blue lines connecting scattered points represent longitudinal scans of the same child. The adjusted value denotes the measure of interest corrected for sex, head motion, and random age effects. yrs, years; LPFC, lateral prefrontal cortex; SPL, superior parietal lobule; IPL, inferior parietal lobule; MPFC, medial prefrontal cortex; VPFC, ventral prefrontal cortex.

At the system level, each subgroup followed a similar system-dependent pattern of nodal entropy (ANOVA, all *p*s < 0.0001 for nine subgroups) (Figure 4A). Specifically, the ventral attention and somatomotor regions showed greater module overlap, while the visual and default-mode regions showed lower module overlap. Quantitative analysis revealed that the entropy of the dorsal attention system significantly increased with age (linear model, *t* = 2.44, *p* = 0.015), while the entropy in the subcortical area significantly decreased with age (linear model, *t* = -3.13, *p* = 0.0019, Figure 4B).

**Figure 4.**
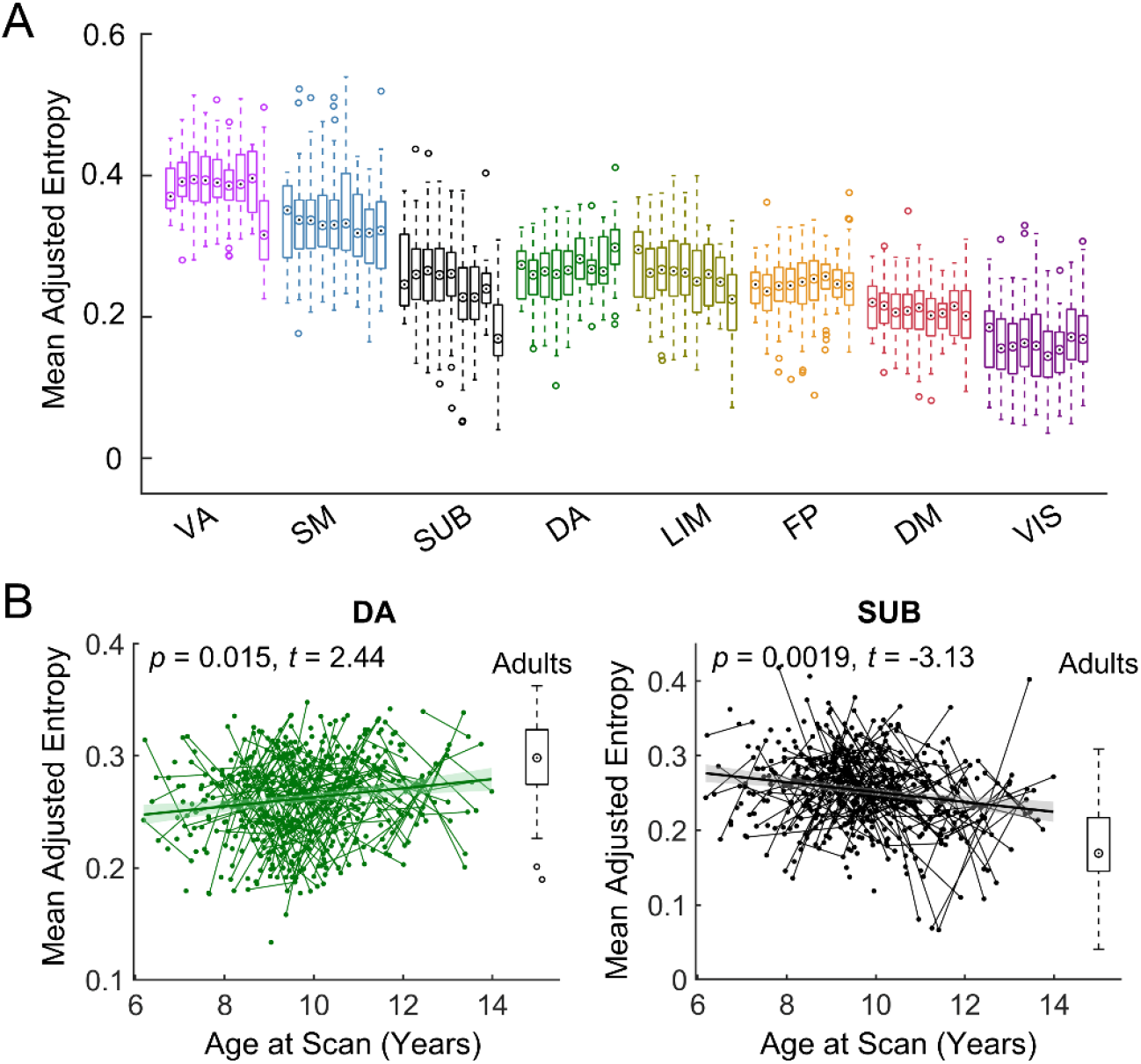
Longitudinal development of the brain functional module overlaps at the system level. **(A)** Distribution of the module overlap within each functional system for each child subgroup and for the adult group. Given a prior system, the bar chart shows the distribution of the average entropy of this system across individuals for eight child subgroups with a one-year interval and the adult cohort. Here, the circle with a dot denotes the median, and the box denotes the interquartile range. **(B)** Two functional systems showing significant developmental changes in functional module overlap. In (B), boxplots represent the distribution of the adult group for reference. Short lines connecting scattered points represent longitudinal scans of the same child. The adjusted value denotes the measure of interest corrected for sex, head motion, and random age effects. VIS, visual; SM, somatomotor; DA, dorsal attention; VA, ventral attention; LIM, limbic; FP, frontoparietal; DM, default-mode; SUB, subcortical.

### Association of cognitive functions with developmental changes in overlapping modules

We next performed a meta-analysis using the NeuroSynth database [49] to explore the potential cognitive significance associated with the developmental changes in network nodes in the overlapping modules. The statistical map of the developmental changes in nodal entropy was divided into 10 bins with decreasing age-related *T* values. We found that the nodal regions that increased with age (bins 0–10 and 10–20), such as the lateral prefrontal cortex, superior parietal lobule, and inferior parietal lobule, were mainly involved in the cognitive terms “motor imagery”, “visual perception”, “spatial”, and “eye movements” (Figure 5). Nodal regions showing decreases with age (bins 80–90 and 90–100), such as the ventral and medial prefrontal cortex and the putamen, were mainly associated with the cognitive terms “speech”, “word form”, “sound”, and “auditory”.

**Figure 5.**
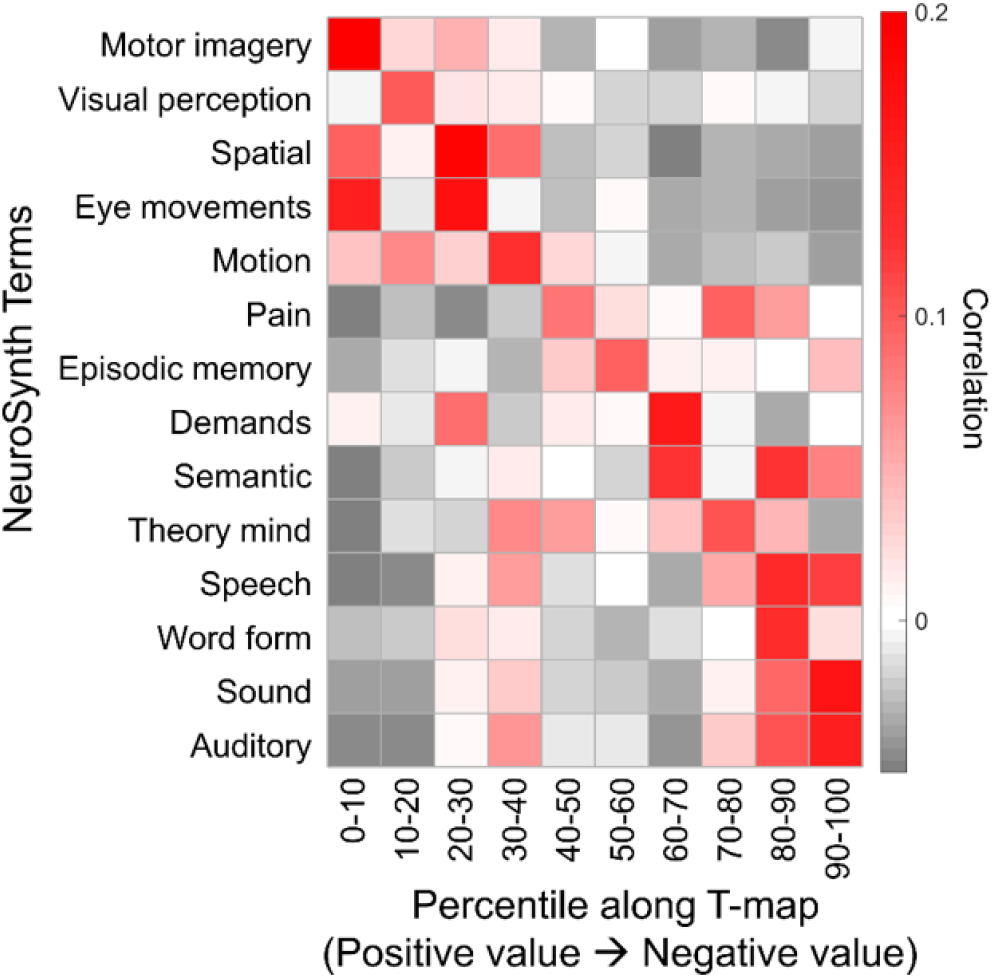
Cognitive function decoding of brain regions. These nodal regions were sorted in descending order according to the developmental changes in the functional module overlap. The results were obtained based on the information in the NeuroSynth meta-analytic database [49].

### Predicting brain age from spatial topography of nodal overlap

We investigated whether the spatial topography of nodal overlap in the network modules could be used to predict individual chronological age. Linear support vector regression (SVR) was used with tenfold cross-validation (Figure 6A). We found that the spatial patterns of nodal entropy significantly predicted individual chronological age (*r* = 0.37, *pperm* < 0.0001; Figure 6B). Regions with high contributions were primarily located in the dorsal attention and default-mode systems (Figure 6C). Furthermore, we found that the nodal contribution weights showed a significant positive correlation with the developmental changes in nodal entropy in terms of age-related *t* values (Pearson’s correlation *r* = 0.46, *p*spin < 0.0001; Figure 6D). The significance level of the spatial similarity was assessed by comparing the observed value to a null distribution. This null distribution was created through 10,000 permutations, generating surrogate maps that maintained the spatial autocorrelation characteristics of the original map [50]. This result suggests that brain regions showing age-related changes play a crucial role in predicting chronological age.

**Figure 6.**
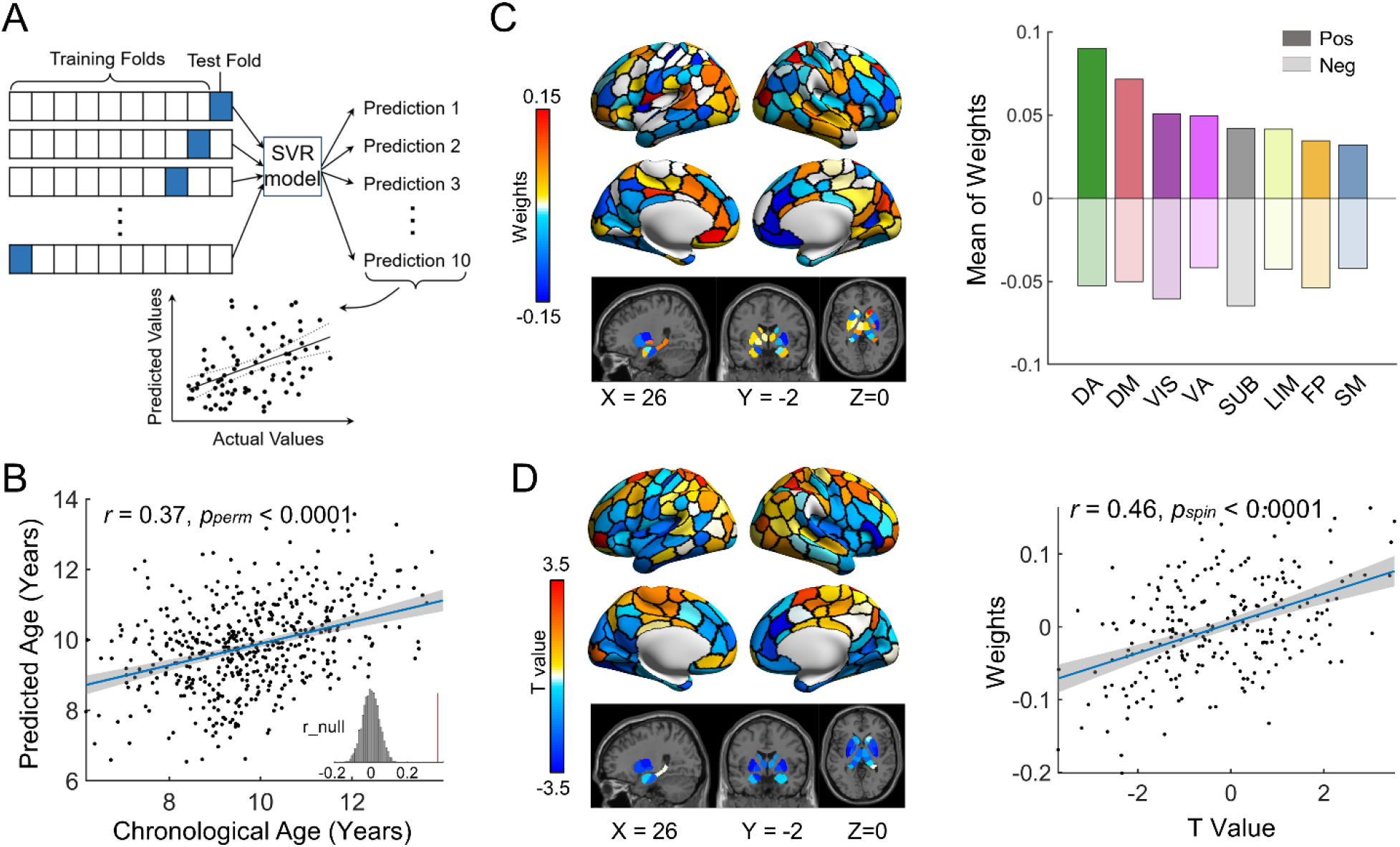
Brain age prediction based on spatial patterns of nodal overlap. **(A)** Schematic representation of the SVR prediction model based on 10-fold cross-validation. **(B)** Accuracy for age prediction using the tenfold SVR model. The gray frequency polygon in the inset displays the null distribution of prediction accuracies based on the permutation tests (*n* = 10,000) by shuffling the actual ages across scans. The red line denotes the actual *r* value. **(C)** Spatial distributions of the nodal contribution in the prediction model. Left: Contribution weight at the regional level. Right: Contribution weight at the system level. Positive and negative weights were separately averaged within each system. **(D)** Left: Spatial pattern of the effect of age on nodal overlap in terms of *T* values. Right: Nodal contribution weight shows a significant positive correlation with the development of nodal overlap based on Pearson’s correlation analysis. Pos, positive; Neg, negative; VIS, visual; SM, somatomotor; DA, dorsal attention; VA, ventral attention; LIM, limbic; FP, frontoparietal; DM, default-mode; SUB, subcortical; SVR, support vector regression.

### Predicting individual nodal patterns of overlapping functional modules from structural brain features

We finally investigated whether the structural features were related to the overlapping modules in children. In this analysis, we included 446 high-quality rsfMRI and dMRI scans from 279 children (aged 6–14 years, F/M = 138/141), with three repeated scans from 42 children, two repeated scans from 83 children and one scan from 154 children. For each brain node of each child, we obtained five morphological measurements (cortical volume, thickness, curvature, folding index, and surface area) using structural MRI data and the fractional anisotropy strength in the white matter networks using diffusion MRI data. For each child, the linear SVR model was used to predict the individual spatial pattern of nodal entropy in the overlapping functional modules by integrating all structural brain features. Figures 7A and 7B show the prediction features and accuracy of a representative child’s scan. We found that the structural features significantly predicted the individual spatial patterns of nodal entropy for 95% of the scans (424/446), with a significance level of *pperm* < 0.05. The prediction accuracy varied across scans (mean ± SD: 0.35 ± 0.10), while the maximum prediction accuracy reached 0.59 (Figure 7C). The prediction contributions varied across structural features, with cortical thickness showing the greatest contribution (Figure 7D).

**Figure 7.**
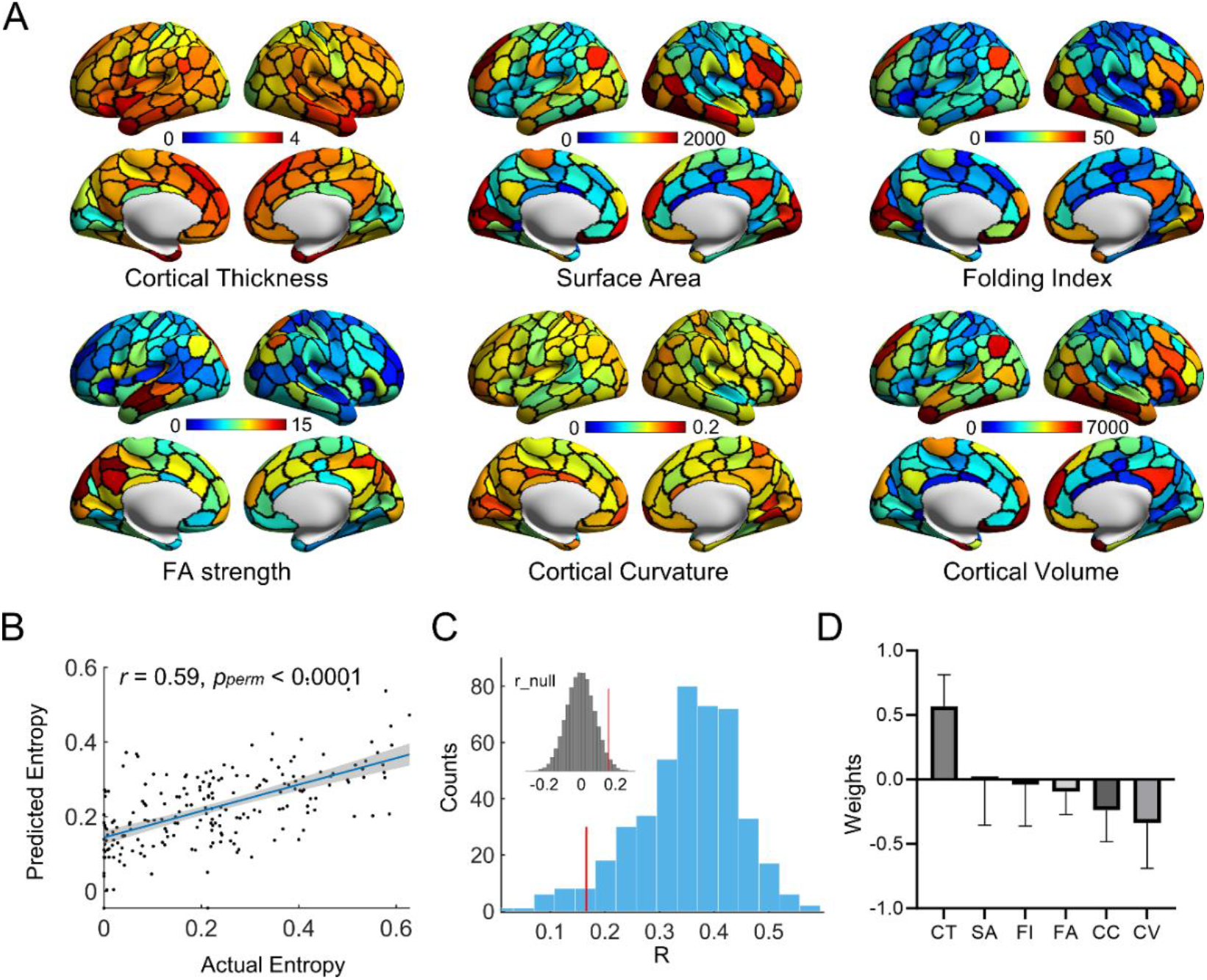
Prediction of individual spatial patterns of nodal module overlap from structural brain features in children. **(A)** Spatial distributions of anatomical features for a representative child’s scan. **(B)** Accuracy of nodal entropy prediction for a representative child’s scan. In (A) and (B), the representative scan was selected as the scan that showed the highest prediction accuracy for the individual map of nodal module overlap. **(C)** Frequency polygon for prediction accuracies of all rsfMRI scans. The inset in the upper left corner denotes the null distribution of the prediction accuracy based on permutation tests. The red line denotes the 95% significance level in the null distribution. **(D)** Prediction contribution of different structural features in the prediction model of all rsfMRI scans. This histogram displays the mean values of anatomical features across subjects, with each bar representing the average value accompanied by its standard deviation. CV, cortical volume; CT, cortical thickness; CC, cortical curvature; FI, folding index; SA, surface area; FA, fractional anisotropy.

### Validation results

We further assessed the reliability of the procedure used for the estimation of nodal overlap in the functional module affiliations. Specifically, we re-estimated the nodal overlap in module affiliations in the adult group by considering the influence of several network construction and analysis strategies, including i) spatial resolution of the parcellation (i.e., Schaefer-100) (Figure S1), ii) network thresholding strategies (i.e., 10% and 20%) (Figure S2A and S2B), iii) module detection algorithms (Figure S2C), and iv) measures for nodal overlap estimation (i.e., involved number) (Figure S2D). We found that the spatial pattern of nodal entropy remained almost unchanged in different cases, suggesting the reliability of the topography of the overlapping modular architecture.

## Discussion

Using a large longitudinal multimodal MRI dataset, we revealed the development of overlapping functional modular organization and its potential structural substrates during childhood and adolescence. Specifically, the spatial distribution of the overlapping modules in children was highly similar to that in the adult cohort, with the ventral attention, somatomotor, and subcortical regions exhibiting greater overlap and the visual and default-mode regions exhibiting lower overlap. The developmental changes in the nodal overlap in module affiliations showed spatial heterogeneity, with significant decreases in the prefrontal cortex and putamen and significant increases in the parietal lobules. Finally, we found that spatial patterns of nodal overlap could be used to predict individual brain ages and were associated with both morphological and white matter features, particularly cortical thickness. Our findings highlighted the spatially inhomogeneous maturation of the overlapping modular architecture and potential structural substrates during childhood and adolescence, providing novel insights into the neural mechanism underlying individual cognitive development.

The presence of the overlapping functional modular architecture is of great value for understanding the functional organization principles of the human brain [31–35]. This framework provides an intuitive characterization of the functional interactions between modules and the diverse roles of brain regions [51, 52]. Compared with previous overlapping module studies in adults [31–35], our study is the first to uncover overlapping functional modular organization in children and adolescents, extending the understanding of brain developmental principles. We found that each functional module comprised nodal regions involved in at least two prior functional systems, regardless of the age subgroup. These systems contained in the same functional module have usually been found to be located in adjacent functional gradients [44].

Taken together, these findings indicate that the overlapping modular architecture may capture intersystem interactions at similar hierarchical levels. Interestingly, most of the prior functional systems were involved in three or more overlapping modules, except for the visual and limbic systems, suggesting diverse functional roles or functional differentiation within each functional system. The default-mode regions were found to be involved in three overlapping modules (i.e., modules I, IV, and VII), which is in line with the previous identification of three subsystems within the default-mode system in both children [39] and adults [53]. From a dynamic perspective, functional modular organization spontaneously reconfigures on a short time scale (e.g., seconds) with regions switching among modules [21]. The observed spatial overlap of functional modules may be a summary of the dynamic reorganization at a long-term scale. However, the relationship between the overlapping and dynamic modular architectures warrants further investigation.

We further found that the topological distribution of the module overlap in children and adolescents showed an adult-like spatially inhomogeneous pattern, suggesting that the overlapping modular architecture is taking shape in children and adolescents. High nodal overlap was primarily located in the ventral attention and somatomotor regions, which is consistent with the findings of a recent study in adults [34]. The high nodal overlap of the ventral attention system may be attributable to its involvement in multiple general domain categories, as revealed in a meta-analytic study [51], including cognition (e.g., attention), perception (e.g., somesthesis) and action domains (e.g., imagination). During a movie watching task, the overlap level of the ventral attention system increased significantly [34], further indicating that the module overlap of the ventral attention system may capture regional involvement in task transitions. The high nodal overlap in the sensorimotor regions indicates the functional diversity of this system, which is consistent with its involvement in multiple modules (Figure 2E). In addition to the widely known sensorimotor functions [54], these regions may be involved in other cognitive functions or behaviors, such as attention allocation [55], speech perception [56], and emotional regulation [57].

The developmental change in the nodal overlap level (i.e., nodal entropy) showed regional heterogeneity. The decreased nodal overlap of subcortical areas may be explained by the gradually decreased involvement of the subcortical system in modules III and V, which are related to somatomotor, attention, and visual functions (Figure 2E). A recent study reported that the strength of cortico-subcortical functional connectivity varies with age, with increasing connections between subcortical and association regions and weakening connections between subcortical and primary regions [58]. The observed age-related decreased overlap of the subcortical regions suggested that the enhanced functional segregation between the subcortical area and the primary system may be more dominant. In the prefrontal cortex, we observed dissociable age-related changes in the extent of nodal overlap for two regions, including the lateral prefrontal cortex and the medial prefrontal cortex. This may be attributed to the fact that these two regions belong to different functional subsystems, i.e., the lateral prefrontal cortex is strongly connected with the default-mode system that is preferentially involved in the regulation of introspective processes, and the medial prefrontal cortex is strongly connected with the dorsal attention system that is mainly involved in the regulation of visuospatial perceptual attention [59]. The significant prediction of an individual’s chronological age based on nodal overlap patterns further demonstrated substantial changes in the overlapping modular architecture during childhood and adolescence.

The functional activity of the human brain is supported and sculpted by the underlying anatomical structure [60]. During childhood and adolescence, both gray matter and white matter of the brain undergo elaborate reconfiguration. The cortical surface area gradually reaches a peak, and the gray matter volume of the whole brain decreases [9]. Moreover, the white matter volume continues to increase [9, 61]. Here, we found that these developing anatomical features of gray matter and white matter could be used to significantly predict the spatial patterns of functional module overlap at the individual level. Of all measures considered, cortical thickness was found to make the greatest contribution. This may be attributed to the remarkable thinning of cortical thickness during childhood and adolescence [9, 62, 63]. Specifically, a previous study has shown that the coordinated development of cortical thickness during adolescence shows a similar pattern of functional connectivity [64]. These results bridge a link between the overlapping functional modular architecture and macroscopic anatomical features in the brain. The potential association with microscale properties (e.g., synaptic pruning and myelination) needs to be further illuminated.

Several issues need to be further considered. First, inner-scan head motion introduces a spatially inhomogeneous bias in functional connectivity estimation, which is age dependent [65]. To reduce the potential influence of head motion, we included 24 head parameters, global brain signals, and “bad” time points during nuisance regression. The individual head motion parameters (i.e., mFD) were also included in the mixed effect model when assessing the age effects. Nevertheless, the potential influence of head movement may not have been completely eliminated. Second, this study used a large longitudinal dataset of children and adolescents, but the age range was limited to school-age children. Future studies should extend the developmental period of interest by adding other datasets from early childhood (even the fetal period) and late adolescence to chart a more complete developmental trajectory from infancy to adulthood. Third, the structure‒function association was established with SVR prediction analysis. It is still unknown how the overlapping functional modular architecture emerges from anatomical features. Other computational models, such as network communication models [66] and large-scale dynamic modeling [67], may be employed to provide the mechanism underlying the brain structure–function relationship. Fourth, we explored the typical developmental changes in the overlapping modules in children and adolescents. Since adolescence is the most common period for the onset of mental disorders [14], the overlapping modular organization may be altered in individuals with neurodevelopmental disorders, such as autism spectrum disorders or attention deficit hyperactivity disorder. Exploring the overlapping modular organization in these disorders may improve the understanding of the pathological mechanism underlying atypical development. Finally, we employed the edge-centric module detection algorithm to detect the overlapping modules, which does not require a prior assumption and is intuitive for understanding multiple module affiliations of nodes. However, until now, there has been no gold standard for evaluating the quality of the detected overlapping functional modular structure in the human brain. The potential biological significance of the overlapping functional modular architecture warrants further research.

## Methods

### Participants

A longitudinal multimodal MRI dataset of 360 healthy children was obtained from the Children School Functions and Brain Development Project (Beijing Cohort) [38, 39]. All participants were cognitively normal, did not use psychotropic medication or had no history of severe traumatic brain injury. These children underwent longitudinal rsfMRI and dMRI scans at intervals of approximately one year. We performed strict quality control on the rsfMRI data and excluded scans with field map errors, excessive head motion (see “Data Preprocessing”), excessive “bad” time points, or T1 artifacts. Finally, 491 scans of 305 participants (aged 6–14 years, F/M = 143/162) remained, including three scans from 47 subjects, two scans from 92 subjects, and one scan from 166 children. Notably, 45 rsfMRI scans were further excluded from the subsequent structural association analysis due to the poor quality of the corresponding dMRI data (see “Data Preprocessing”). In addition, 446 rsfMRI scans from 279 subjects (aged 6–14 years, F/M = 138/141) were used in the structure‒function association analysis, including three scans from 42 subjects, two scans from 83 subjects and one scan from 154 subjects. In addition, we employed rsfMRI scans of 61 healthy young adults (aged 18–29 years, F/M = 37/24) for comparison. All participants or their parents/guardians provided written informed consent, and this study was approved by the Ethics Committee of Beijing Normal University.

### Imaging acquisition

Multimodal magnetic resonance images were acquired on a 3T Siemens Prisma scanner with a 64-channel head coil at the Center for Magnetic Resonance Imaging Research at Peking University. Both children and adults underwent multimodal scanning using the following protocols.

*(i) Functional MRI.* Resting-state scans were acquired using an echo-planar imaging sequence: repetition time (TR) = 2000 ms, echo time (TE) = 30 ms, flip angle = 90°, field of view (FOV) = 224 × 224 mm^2^, acquisition matrix = 64 × 64, slice number = 33, and slice thickness/gap = 3.5/0.7 mm. All participants were asked to fixate on a bright crosshair displayed in the center of the scanner screen. The total duration of the rsfMRI scans was eight minutes (i.e., 240 volumes).
*(ii) Field maps for functional MRI.* The scans were acquired using a 2D dual gradient-echo sequence: TR = 400 ms, TE1 = 4.92 ms, TE2 = 7.38 ms, flip angle = 60°, FOV = 224 × 224 mm^2^, acquisition matrix = 64 × 64, slice number = 33, and slice thickness/gap = 3.5/0.7 mm.
*(iii) T1-weighted structural MRI.* The scans were acquired using a sagittal 3D magnetization prepared rapid acquisition gradient-echo (MPRAGE) sequence: TR = 2530 ms, TE = 2.98 ms, flip angle = 7°, FOV = 256 × 224 mm^2^, acquisition matrix = 256 × 224, inversion time = 1100 ms, slice number = 192, slice thickness = 1 mm, and bandwidth = 240 Hz/Px.
*(iv) Diffusion MRI.* The scans were acquired using a high angular resolution diffusion imaging (HARDI) sequence: TR = 7500 ms, TE = 64 ms, flip angle = 90°, FOV = 224 × 224 mm^2^, acquisition matrix = 112 × 112, slice number = 70, slice thickness = 2 mm, and bandwidth = 2030 Hz/Px. The complete sequence consisted of 64 diffusion-weighted directions (b-value = 1000 s/mm^2^) and 10 nondiffusion-weighted directions (b-value = 0 s/mm^2^).

### Data preprocessing

*(i) Functional MRI data.* The functional images of all the children were preprocessed using SPM12 (https://www.fil.ion.ucl.ac.uk/spm) and DPABI 3.0 [68]. First, we removed the first 10 volumes and performed slice-timing correction. Next, we applied field map correction to reduce geometric distortion and realigned the volumes over time. After realignment, 94 rsfMRI scans were excluded due to excessive head motion with a criterion of maximum head motion > 3 mm or 3° or mean framewise displacement (mFD) [69] > 0.5 mm. Then, the functional images were coregistered with individual T1 images and spatially normalized to a custom pediatric template using a unified segmentation algorithm [70] with the following steps: i) individual structural images were initially segmented into three tissue (i.e., gray matter, white matter, and cerebrospinal fluid) probability maps by using the Chinese Pediatric Atlas (CHN-PD) (6–12 years) [71] as a reference; ii) the resulting spatially normalized maps for each tissue type were averaged across scans to generate the custom tissue templates; iii) individual structural images were segmented again using the custom tissue templates as a reference; and iv) the functional images were spatially normalized using the transformation parameters estimated from the second segmentation of structural images. For the child cohort, custom tissue maps were used to improve the accuracy of spatial normalization. The normalized functional images were then resampled into 3-mm isotropic voxels and underwent spatial smoothing using a Gaussian kernel (full-width at half-maximum = 4 mm), linear detrending, and nuisance signal regression. A series of regressors were included in the nuisance regression, including 24 head motion parameters [72], “bad” time points with FD above 0.5 mm, white matter signals, cerebrospinal fluid signals, and global brain signals. Finally, we performed temporally band-pass filtering (0.01-0.1 Hz) on the images. For the adult cohort, the preprocessing procedures were conducted in the same way as for the child cohort, except that the functional images were normalized to the Montreal Neurological Institute (MNI) standard space.
*(ii) T1-weighted MRI data.* The T1-weighted images were preprocessed using FreeSurfer v6.0 [73]. First, we performed intensity normalization and removed nonbrain tissue using the HD-BET algorithm [74], during which the automatically extracted brain tissue maps replaced the default maps (i.e., “brainmask.mgz”) in FreeSurfer to improve accuracy. Next, we conducted tissue segmentation and cortical reconstruction on individual T1 images. Notably, the longitudinal processing stream of FreeSurfer was selected to obtain robust and reliable morphological measurements [75]. A trained researcher visually inspected the cortical reconstruction results to ensure that the correct boundaries were estimated. Finally, we obtained five local gray matter morphological features, including cortical volume, thickness, curvature, folding index, and surface area for each vertex.
*(iii) Diffusion MRI data*. The diffusion images were preprocessed using MRtrix 3.0 [76], FSL 6.0.1 (https://fsl.fmrib.ox.ac.uk/fsl/fslwiki/FSL), and ANTs [77]. First, the diffusion data were denoised, and Gibbs ringing artifacts were removed. Next, we corrected the eddy current-induced distortions, subject movement, and signal dropout for each scan using the FSL eddy tool [78]. Notably, 45 rsfMRI scans were further excluded from the structure‒function association analysis due to missing images, excessive head motion (maximal motion > 3 mm), or considerable signal dropout in the corresponding diffusion images. Then, field map correction was employed to reduce susceptibility by using FSL (epi_reg script). Finally, B1 field inhomogeneity was corrected with the N4 algorithm [79].

### Identification of the overlapping modular architecture

We identified the overlapping modular architecture in the brain functional networks by employing edge-centric module detection [40] (Figure 1B). Briefly, we first constructed a traditional functional network comprising nodal regions and interregional connectivities (i.e., edges). Then, we constructed the corresponding weighted edge graph representing the similarity between edges. Finally, we identified the module affiliations of each nodal region according to the module assignments of its edges in the corresponding edge graph.

#### Functional network construction

We constructed a brain functional network comprising 232 nodal regions for each rsfMRI scan of each participant. In this functional network, cortical nodes were defined based on a recently developed functional parcellation comprising 200 cortical regions (i.e., Schaefer-200) [80], and subcortical nodes were defined according to a subcortical functional parcellation comprising 32 regions [43]. We extracted the mean time series for each nodal region and estimated Pearson’s correlation coefficients between every pair of nodes. Then, a weighted functional network was obtained by thresholding the correlation matrix with a density of 15% (i.e., 4020 edges) to exclude potential spurious or weak correlations. Negative correlations were not considered due to their controversial physiological interpretations [81, 82].

#### Edge graph construction

For each functional network, we constructed a corresponding weighted edge graph that denoted the similarity between edges (Figure 1B). For simplicity, we considered only directly connected edges that shared at least one common node, and the similarity between edges without common nodes was assumed to be zero. The Tanimoto coefficient [30, 83] was used to incorporate the edge weight information. For a pair of edges *eik* and *ejk* that share a common node *k*, their similarity was defined based on the similarity in the connection profiles between nodes *i* and *j*:

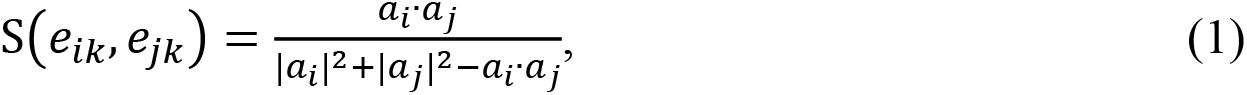

where *a*_*i*_ · *a*_*j*_ represents the dot product of the two vectors *a*_*i*_ and *a*_*j*_, and *a*_*i*_ = (*Ã*_*i*1_, *Ã*_*i*2_, … , *Ã*_*iN*_) represents a modified connectivity profile of node *i*. Specifically, each element *Ã*_*ij*_ is defined as

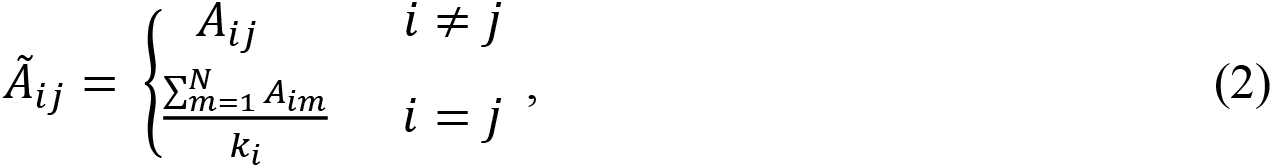

where *A*_*ij*_ is the functional connection strength between nodes *i* and *j* and *ki* is the number of edges of node *i*. Of note, nonzero diagonal elements were included in Eq. (2) to make the similarity definition, i.e., Eq. (1), applicable to extreme cases in which nodes *i* and *j* are directly connected.

### Identifying the overlapping modular architecture based on edge-centric module detection

Given a functional network of interest, we first detected the modular structure in the corresponding edge graph and then determined the module affiliations of each nodal region according to the module affiliations of its edges. Here, the Louvain algorithm [41] was employed to detect the modular architecture in the large-scale edge graph, and each edge was assigned to a specific module. Each nodal region was then assigned to one or more modules due to diverse module affiliations of its edges, leading to spatial overlap between the functional modules. A nodal region whose edges were involved in two or more modules was defined as an overlapping region. Notably, the detected module partition in the edge graph varied slightly across each instance of detection due to the heuristic property of the Louvain algorithm. The module number of the functional network was defined as the module number that appeared most frequently among 100 instances, and the other measurements regarding the overlapping modular structure were taken as the average across 100 instances of identification.

### Topography analysis of the overlapping modular architecture at the group level

To illustrate the spatial patterns of the overlapping modular structure at different ages, we detected the modular architecture at the group level. All the children’s rsfMRI scans were divided into eight subgroups with one-year intervals, and the adult group was set as the ninth subgroup for comparison. A group-level weighted functional network was constructed for each subgroup by averaging individual functional correlation matrices followed by network thresholding (i.e., density = 15%). Then, we detected the modular structure in the corresponding edge graph for each subgroup with 100 instances of module detection. To obtain a stable module division, we conducted the following two steps: i) computed the module co-occurrence matrix between each pair of edges [84], wherein each element denoted the proportion of instances in which a pair of edges were assigned to the same module; and ii) applied the modular detection algorithm to the module co-occurrence matri× 100 times. We iterated these two steps until the module partition remained unchanged across multiple instances. The final version of the module partition of edges was used to infer the overlapping modular architecture of the functional network.

Considering the spatial overlap of functional modules, we obtained a spatial map for each functional module separately. Nodal values in the map denote the proportion of edges assigned to this module. To compare the modules observed in each child subgroup with those in adults, we further matched the functional module maps between the adult subgroup and all the child subgroups. Given a module map of interest in the adult group, a matching module map was selected as the map showing the maximal similarity for each child subgroup. The spatial similarity between two maps was calculated as the Pearson’s correlation coefficient across nodal regions.

To further assess the functional system dependence of the overlapping modular structure, we mapped each cortical or subcortical node to one of the prior functional systems, including the visual, somatomotor, dorsal attention, ventral attention, limbic, frontoparietal, and default-mode systems [42] and the subcortical area [43]. For each module, we quantified its spatial pattern by estimating the percentage of nodes distributed in each functional system.

### Measure of the extent of module overlap in brain regions

Given an overlapping modular architecture, a node may be involved in multiple modules due to the diverse module affiliations of its edges. Here, entropy was used to measure the extent of nodal overlap by quantifying the distribution of module affiliations of edges attached to a node (Figure 1B). Given a node *i*, the entropy [34] was defined as

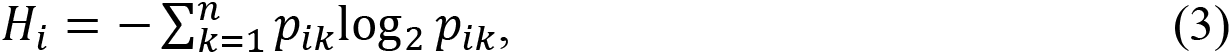

where *n* is the number of modules involved with this node, *pik* represents the proportion of its edges participating in module *k*, and 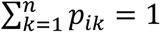. Nodal entropy was further normalized to a range of 0 to 1 by dividing it by log2*n* [34]. A higher normalized entropy value indicates greater overlap, suggesting that the distribution of edges among different modules is more homogeneous and diverse. For each scan of each participant, we obtained a nodal module overlap map in terms of nodal entropy.

To illustrate the patterns of nodal overlap at different ages, a group-level entropy map was separately generated for each child subgroup and the adult group by averaging individual entropy maps within the subgroup. Then, we estimated the spatial similarity of the entropy maps between each of the child subgroups and the adult subgroup by calculating Pearson’s correlation coefficients across nodal regions. We further employed one-way analysis of variance (ANOVA) models to explore whether the nodal overlap level was functional system-dependent for each subgroup. The functional systems were defined based on the prior seven functional systems [42] and the subcortical area [43].

### Analysis of developmental changes in the overlapping modular architecture

To explore the developmental changes in the overlapping modular architecture, we assessed the effects of age on a series of brain measures, including the modular number, the modularity of the edge graph, and the nodal overlap level, in terms of entropy values at the system and nodal levels. For each brain measure of interest, the age effects were estimated by using a mixed effect model [47, 48]. The mixed model is suitable for a longitudinal dataset with irregular intervals between measurements and is applicable to cases of missing time points. The model parameters were estimated with the maximum likelihood method. Considering the potential linear and quadratic age effects, we employed both the linear model and the quadratic model of the age effects. The optimal model was selected for each brain measure based on the Akaike information criterion (AIC) [85]. Sex and the in-scanner head motion parameter (i.e., mFD) were included as covariates within each model.

The linear model was defined as

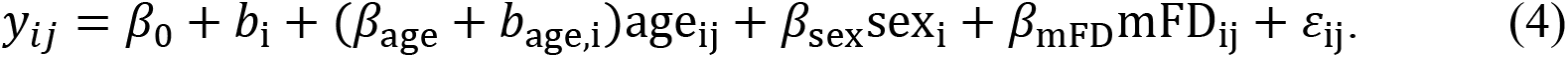

The quadratic model was defined as

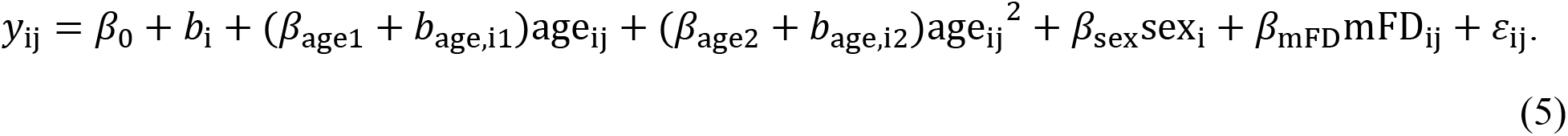

In these equations, *yij* represents the brain measure of subject *i* at the *j*th scan, *β*_*age*_ represents the fixed effect, *bage,i* represents the random effect, and *ɛ*_*ij*_ represents the residual. For the nodal and system-level analyses, the significance level of the results was corrected for multiple comparisons across nodes using the false discovery rate (FDR) method [86].

### Associations between cognitive functions and developmental changes in nodal overlap

To explore the cognitive significance of developmental changes in nodal overlap, we performed a meta-analysis using the NeuroSynth database (www.neurosynth.org) [49]. We first sorted the developmental changes in nodal entropy (i.e., *t* values) in decreasing order and then divided the brain map into 10 bins (i.e., 10% apart as a box). Each bin was binarized to generate a brain mask. Next, we calculated the Pearson’s correlations between each mask and all the cognitive term maps available in the database. We selected the top two associated cognitive terms for each mask and removed six overlapping terms among the 10 masks. Finally, 14 cognitive terms were used to depict the distribution of cognitive functions across different levels of developmental changes.

### Predicting brain age from spatial patterns of nodal overlap

We used linear support vector regression (SVR) with tenfold cross-validation to test whether individual spatial patterns of nodal overlap could be used to predict a participant’s chronological age. The nodal entropy map was set as features for each participant. First, all the entropy maps were divided into ten subsets with similar age distributions. This division strategy is better than the random splitting method, which can reduce the sampling bias among the subsets [87, 88]. Of the ten subsets, one was designated as the testing set, and the remaining nine subsets were used as the training set. Next, we linearly scaled each feature across individuals between zero and one in the training set and applied the estimated scaling parameters to the testing set. Then, we trained the prediction model for individual chronological ages based on the training set. Finally, we quantified the prediction accuracy by calculating the Pearson’s correlation coefficients between the predicted scores (i.e., ages) obtained from the SVR model and the actual scores (i.e., chronological age). To assess the statistical significance of the prediction accuracy, we generated a null distribution of accuracy based on permutation tests (n = 10,000) by shuffling the actual scores across scans. To reduce the influence of confounding factors, we further corrected for sex, in-scanner head motion (i.e., mFD), and random age effects from the nodal entropy values prior to the prediction analysis. To determine the contribution of nodal features to the prediction model, we trained another SVR model using all the participants to improve the estimation accuracy [87, 88]. The resulting regression coefficients were regarded as the weights denoting the importance of all features. To further clarify the system dependence of the weights, we also classified the nodal weights into eight functional systems, including seven cortical functional systems [42] and the subcortical area [43]. The positive and negative weights were separately averaged within each system. Here, the SVR model was implemented using the LIBSVM toolbox in MATLAB with the default parameters (https://www.csie.ntu.edu.tw/~cjlin/libsvm/) [89].

### Analysis of nodal integrity in the white matter structural network

In addition to local morphological measures, we also considered the potential influence of nodal integrity in the white matter structural network. Structural networks were generated from the preprocessed diffusion images using DSI Studio (http://dsi-studio.labsolver. org). First, we tracked the fasciculus and obtained diffusion anisotropy parameters (i.e., fractional anisotropy, FA). Specifically, we employed the generalized q-sampling imaging (GQI) algorithm [90] with a diffusion sampling length ratio of 1.25 for deterministic tractography. The Otsu threshold was 0.6. The tracking procedure was terminated if the turning angle was > 45° or if the fibers reached the borders of the cerebrospinal fluid or subcortical areas. Ten million streamlines were generated with a step size of 0.625 mm, and only tracts with a length between 6∼250 mm were retained for subsequent analysis. Next, the reconstructed streamlines were projected to the Schaefer-200 atlas, which has been registered to the native space. Finally, we calculated the FA values between every pair of nodes (i.e., the mean FA values along all reconstructed streamlines between two nodes) and further summed the FA values between a node and all the other nodes. This metric was defined as nodal FA strength, which indicates the microstructural integrity of the fiber bundles attached to a node [91].

### Predicting individual nodal overlap maps from structural features

To evaluate whether the spatial pattern of the nodal overlap maps was associated with anatomical architecture, we performed a prediction analysis based on the SVR model for each scan of each participant. We considered five morphological measurements (cortical volume, thickness, curvature, folding index, and surface area) in the gray matter and the FA strength in the white matter. For each node, morphological features were extracted as the average value within the nodal region based on the Schaefer-200 atlas, which has been registered to the native space. The prediction was conducted using the same framework as that used in the age prediction mentioned above. Tenfold cross-validation was also used here. In each validation instance, ten percent of the nodal regions were designated as the testing set, and the remaining nodes were set as the training set. We predicted nodal entropy values for each scan by integrating anatomical features from both gray matter and white matter. To assess the statistical significance of the prediction accuracy, we generated a null distribution of accuracy using permutation tests (*n* = 100) by shuffling the actual entropy across nodes for each scan, thus leading to 100 × 446 (scans) permutation instances in total. The 95% significance level was determined according to the null distribution containing all 44,600 instances. To assess the prediction contribution of anatomical features, we trained another SVR model for each scan using all the nodal regions in the whole brain to improve estimation accuracy [87, 88]. The resulting regression coefficients were regarded as the weights denoting the importance of all features.

### Validation analyses

To ensure the reliability and robustness of the topography of the overlapping modular architecture, we investigated the potential influence of several network construction and analysis strategies in the adult cohort. Specifically, we examined the potential influence of functional parcellation on the nodal definition, network thresholding strategy, edge module detection algorithm, and nodal overlap estimation. First, we analyzed the spatial resolution of the functional parcellation. For the main analysis, we constructed a brain functional network comprising 200 cortical regions [80] and 32 subcortical regions [43]. To further validate the influence of spatial resolution, we reconstructed whole-brain functional networks, during which the cortical nodes were defined based on the Scheafer-100 atlas, which comprises 100 cortical regions [80]. Second, we examined the network thresholding density by obtaining weighted functional networks using two other network densities (i.e., 10% and 20%). Third, we explored the influence of the module detection algorithm. In addition to the Louvain algorithm [41], we employed the eigenspectral analysis method [92] to detect the modular structure of the edge graph. Finally, we quantified the nodal overlap level by considering the number of modules involved, which provides a more intuitive understanding. For each nodal region, we calculated the involved module number as the number of modules involved in its edges. The larger the involved number, the higher the nodal overlap in module affiliations.

## Author contributions

**Conceptualization:** Tianyuan Lei, Xuhong Liao, and Yong He.

**Data curation:** Tianyuan Lei, Xiaodan Chen, Weiwei Men, Yanpei Wang, Leilei Ma, Ningyu Liu, Jing Lu, Gai Zhao, Yuyin Ding, Yao Deng, Jiali Wang, Rui Chen, Haibo Zhang, Shuping Tan, Jia-Hong Gao, Shaozheng Qin, Sha Tao, Qi Dong, and Yong He.

**Formal analysis:** Tianyuan Lei and Xuhong Liao.

**Funding acquisition:** Qi Dong, Sha Tao, Xuhong Liao and Yong He.

**Investigation:** Tianyuan Lei, Xinyuan Liang, and Lianglong Sun.

**Methodology:** Tianyuan Lei, Xinyuan Liang, Lianglong Sun, Yunman Xia, Mingrui Xia, Tengda Zhao, Xiaodan Chen and Xuhong Liao.

**Project administration:** Xuhong Liao and Yong He.

**Supervision:** Xuhong Liao and Yong He.

**Visualization:** Tianyuan Lei.

**Writing – original draft:** Tianyuan Lei, Xuhong Liao, and Yong He.

**Writing – review & editing:** Tianyuan Lei, Xuhong Liao, Yong He, Mingrui Xia, and Tengda Zhao.

## Data Availability Statement

All code and data used to perform the analyses are available on GitHub at https://github.com/helab207/Development-of-the-overlapping-modular-structure-in-human-brain-functional-networks.

## Supplementary Information

**Figure S1.**
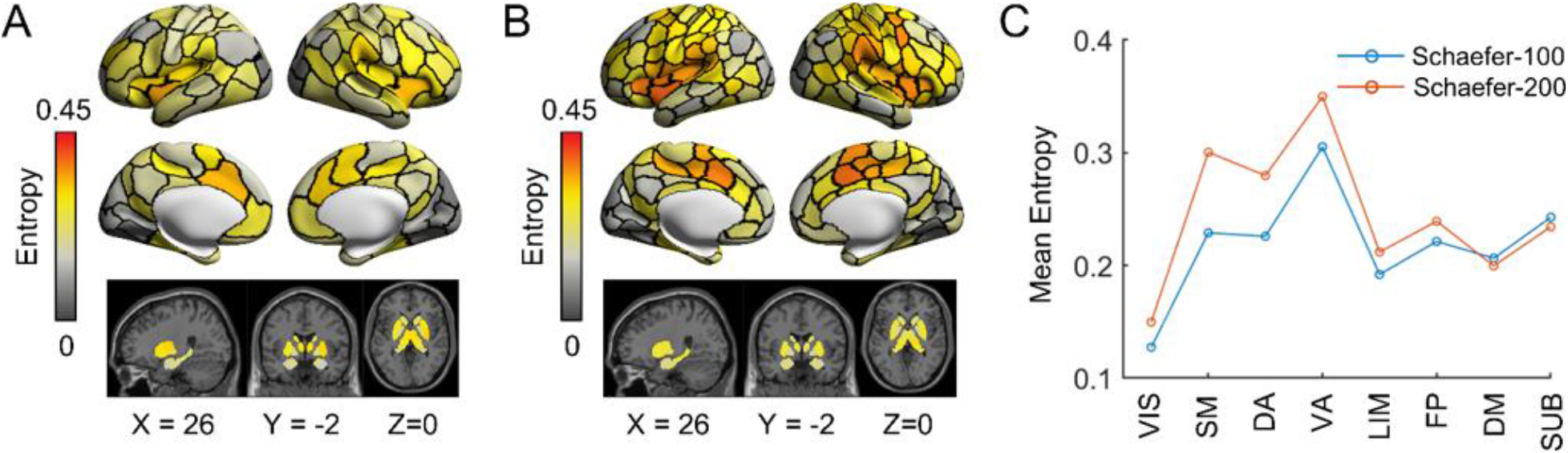
The spatial patterns of the nodal module overlap at different spatial resolutions. For the adult group, the overlapping modular architecture was separately detected in the group-level functional networks obtained from different functional parcellations. **(A)** Nodal overlap in the functional networks with coarse parcellation. This network comprised 100 cortical nodes obtained from the Schaefer-100 atlas (Schaefer et al., 2018) and 32 subcortical regions (Tian et al., 2020). **(B)** Nodal overlap in the functional networks with a fine parcellation (i.e., main results). This network comprised 200 cortical nodes obtained from the Schaefer-200 atlas (Schaefer et al., 2018) and 32 subcortical regions (Tian et al., 2020). **(C)** Extent of nodal overlap for eight systems at different spatial resolutions. Similar distributions of nodal overlap were observed between the two parcellations. VIS, visual; SM, somatomotor; DA, dorsal attention; VA, ventral attention; LIM, limbic; FP, frontoparietal; DM, default-mode; SUB, subcortical.

**Figure S2.**
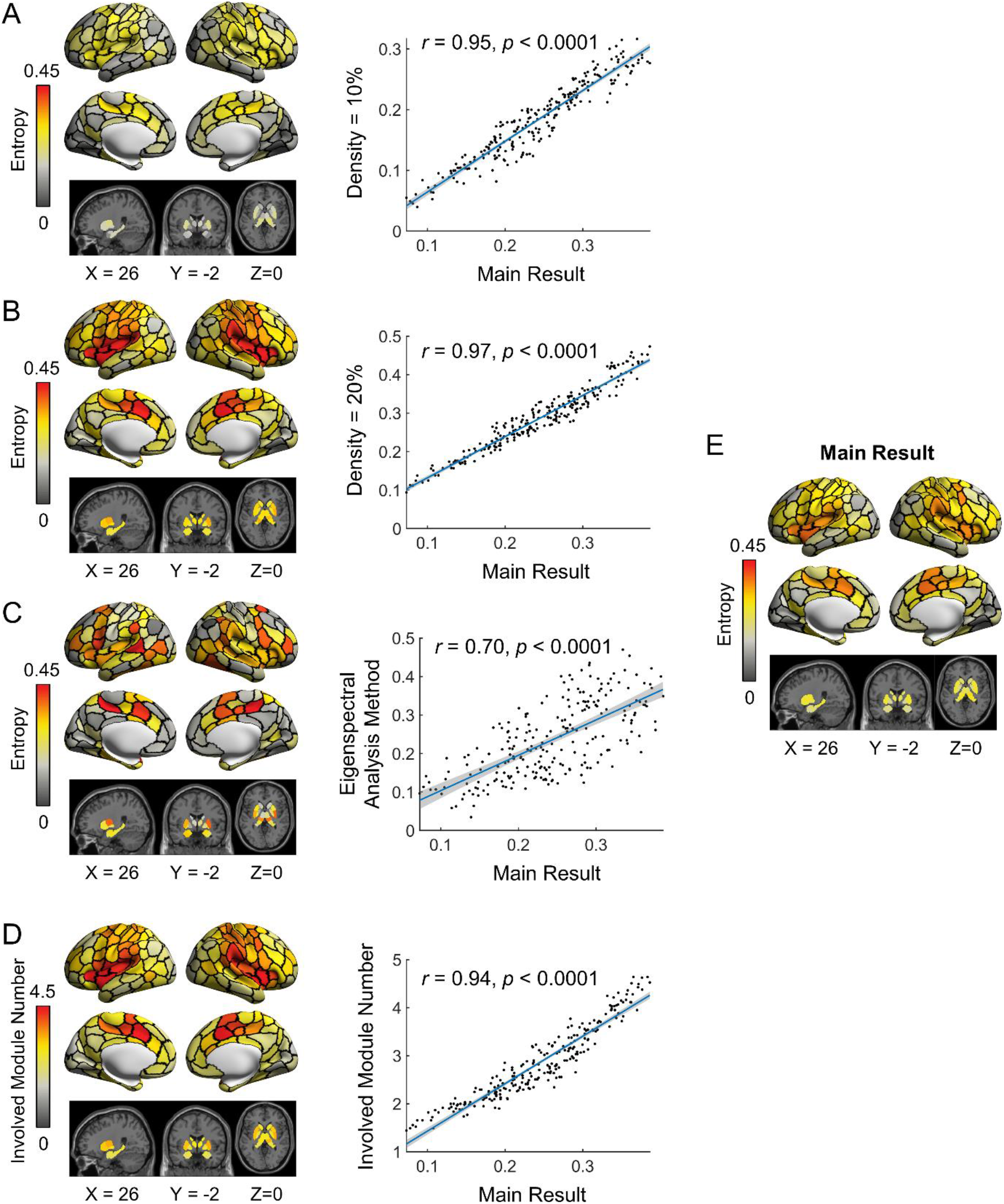
The spatial patterns of the nodal module overlap under different network analysis strategies and their relationships with the main results. **(A)** Network density of 10% for functional network construction. **(B)** Network density of 20% for functional network construction. **(C)** Eigenspectral analysis for module detection in the edge graph. **(D)** The number of involved modules used to quantify the extent of nodal module overlap. **(E)** Main result as a reference. In each case, all the network construction and analysis strategies were set to be the same as those in the main analysis, except for the strategy of interest. All correlations were assessed with Pearson’s correlation across nodal regions.

